# Analytical limits of hybrid identification using genetic markers: an empirical and simulation study in *Hippolais* warblers

**DOI:** 10.1101/026781

**Authors:** JO Engler, S Twietmeyer, J Secondi, O Elle, A Hochkirch

**Affiliations:** Zoological Researchmuseum Alexander Koenig, D-53113 Bonn, Germany; Department of Wildlife Sciences, University of Göttingen, D-37077 Göttingen, Germany; Nationalparkforstamt Eifel, Urftseestraβe 34, 53937 Schleiden-Gemünd; Department of Biology, Molecular Ecology and Evolution Laboratory, Lund University, Ecology Building, 223 62 Lund, Sweden; Biogeography Department, Trier University, D-54286 Trier, Germany

## Abstract

Hybridization is known to occur in a wide range of avian species, yet the rate and persistence of hybridization on populations is often hard to assess. Genotyping using variable genetic marker sets has become a common tool to identify hybrid individuals, however assignment outputs can differ depending on the marker set used. Here, we study hybrid assignment in two sibling *Hippolais* warblers, where hybrid assignment has shown to differ between SSR and AFLP markers. Simulation of heterospecific individuals as well as backcrosses (typed using SSR markers) reveals a rapid loss of assignment probability in higher backcross generations.

However, the characterization of F1 hybrids was clearly distinguished from both parental taxa. The differences in marker sets are not contradictory but complementary. The rate of hybridization is lower than previously expected with AFLP markers but introgression might be long-lasting. This could be either due to differences in power of the marker systems used or due to non-neutral variation covered by AFLP but not SSR markers. We call for more attention to be paid regarding the potential limits of classical marker systems to investigate hybridization and its persistence in natural systems.

## Introduction

Hybridization is known to occur in a wide range of avian species (McCarthy 2006). However, the rate and persistence of hybridization on populations is often hard to assess. Genotyping using variable genetic marker sets has become a common tool to identify hybrid individuals, yet assignment outputs can differ depending on the marker set used. The sibling old world warblers *Hippolais icterina* and *H. polyglotta* share a moving contact zone in Central Europe. Heterospecific breeding has been observed for decades suggesting crosses between both species (e.g. Ferry 1980, Faivre & Ferry 1989). In later studies, hybridization was confirmed by analyzing morphometric (Faivre et al. 1999) and genetic data based on amplified fragment length polymorphism (AFLP) markers (Secondi et al. 2006).

The prominent genetic signal however disappeared completely in a recent reanalysis of the same genetic samples based on microsatellite (SSR) markers (Engler et al. *unpublished*) despite a high potential hybrid detectability in the marker set used (Engler et al. 2014). In a further study comparing genetic changes in three German populations at the range edge over an eight year period, and using additional samples, (Engler et al. 2013) also showed no evidence of hybridization. This questions the true frequency to which hybridization between these species occurs.

Recently, efforts were focused to find hybrids in areas where both species occur in sympatry and prove their mixed ancestry using the same set of SSR markers as used in Engler et al. (*unpublished*) and following the protocol of Engler et al. (2014). In this note, we present three cases of hybridization between the two *Hippolais* warblers.

Based on a simulation of genotypes between both parental species, our aim was to explore how fast hybrid genotypes will diminish in the gene pool of the expansive species and whether the identified hybrid individuals would fall into either direct hybrid or backcross categories. We therefore assessed the likelihood of detecting and differentiate hybrid allele combinations as well as backcrosses under the given marker set and link them to the detected genotypes.

## Methods

### Field work

Searches for potential hybrid individuals were conducted randomly in suitable habitat along the German part (Rhineland-Palatinate) of the contact zone between *Hippolais polyglotta* and *H. icterina* in the breeding seasons between 2008 and 2012. A total of 13 potential hybrid candidates were identified in the field based upon mixed song and/or intermediate wing morphologies. Birds were captured using a mist net and song playback of either *H. polyglotta* or *H. icterina* song. Blood samples (c. 10 μl) were taken from the ulnar vein and the birds were released afterwards. For birds where blood sampling was not possible, one single feather was taken from the breast before release. Additional feather samples were collected from the Netherlands (Region Limburg) by Boena van Noorden from a mixed breeding pair and its offspring (n = 5) in 2009.

### Genotyping

DNA was extracted from feather (stored dry) or blood (stored in ethanol or EDTH buffer solution) samples. Feather DNA extraction was done with the DNEasy Blood and Tissue Kit (Qiagen), whereas DNA from blood samples was extracted using the High Pure PCR Template Preparation Kit (Roche) following the manufacturers protocol. Samples of potential hybrid candidates were genotyped at eight microsatellite loci (*ASE19, ASE34, ASE37, ASE46, TG05_053, TG11_011, ApCo46, Tc.11B4E_2nd*). This set has been identified by Engler et al. (2014) as suitable for hybrid identification and was applied in Engler et al. (submitted) for inter-specific analyses. Different primer pairs were pooled together according to their dye label, annealing temperature, and size into four different sets for multiplex PCR (Table 1). PCR protocol following Engler et al. (submitted).

**Table 1:** Multiplex sets of used microsatellite loci.

| **Marker & Locus**<br>**EMBL accession no.** | **Multiplex**<br>**set** | **T<sub>a</sub>**<br>**[C°]** | **Dye**<br>**label** | **Allele**<br>**size (bp)** |
| --- | --- | --- | --- | --- |
| Ase34<br>*AJ276636* | 1 | 54 | HEX | 204-240 |
| TG05_053<br>*CK314425* | 1 | 54 | 6FAM | 210-214 |
| ASE46<br>*AJ276775* | 2 | 57 | HEX | 240-246 |
| ApCo46-ZEST<br>*AF520885* | 3 | 65 | 6FAM | 213-215 |
| Tc.11B4E-CEST (2nd set)<br>*AF036266* | 3 | 65 | 6FAM | 396-402 |
| TG11_011<br>*CK308096* | 4 | 56 | HEX | 209-213 |
| ASE37<br>*AJ276639* | 4 | 56 | 6FAM | 234-238 |
| ASE19<br>*AJ276376* | 4 | 56 | 6FAM | 170-180 |

Species assignment was conducted using Structure v2.3.3 (Prichard et al. 2000, Falush et al. 2003) with a set of 20 allopatric samples from the respective parental taxa (ten samples from either species). In Structure, we set the number of groups, K = 2 according to the two species used in the data set. Our previous study has shown that there is no intra-specific structuring in either species (Engler et al. submitted), so we are confident that K_max_ = 2. Structure was run with a chain length of 2,000,000 generations and a burn-in length of 500,000 generations. We used an admixture model assuming correlated allele frequencies and kept all other settings in default mode.

### Simulations

We compiled an R-script in R 3.0.2. (R development core team, 2014) that simulated two different scenarios: In the first simulation, we generated 50 different F1 hybrids as well as 25 F2 hybrids and first generation backcrosses with either parental species based on 50 randomly chosen individuals of each parental species. In a second simulation, we used the same individuals from the parental species to simulate five generations of backcrosses with HP. This scenario should highlight the duration (in generations) how long a hybridization signal persists under a situation of range expansion in HP (i.e. mating partners of just one species become available after the hybridization event, Secondi et al. 2006). Individuals were selected from allopatric sites only (*sensu* Secondi et al. 2006, Engler et al. submitted) to account for potential introgression events in samples located nearby the contact zone (Secondi et al. 2006). Only individuals for which all loci were successfully amplified were used for the simulations.

Afterwards, assignment was performed again using STRUCTURE by using the same settings as described above. From this, the average assignment score was calculated with the respective confidence intervals.

## Results & Discussion

### Hybrid candidates

From the 13 potential candidate hybrids sampled in Germany only two could not be clearly assigned to a parental species. From the remaining eleven individuals, eight were assigned as *H. polyglotta* and three as *H. icterina.* The inferred ancestry in STRUCTURE of one of the two individuals that could not be clearly assigned (DT12) was 0.508 for *icterina* and 0.492 for *polyglotta* respectively. The second individual (DT06) was less clear and shows an ancestry to *icterina* of just 0.125 and to *polyglotta* of 0.875. However this ancestry value still is outside the 95% CI of the estimated ancestries of allopatric *polyglotta* samples (mean = 0.946, 95% CI = 0.939- 0.954).

From the mixed breeding pair and its offspring from the Netherlands, a sample from the father was missing. The mother was *icterina*. Interestingly, from the two samples from the offspring, one individual (NL05) was classified as pure *icterina* genotype, whereas its sibling (NL04) showed a hybrid ancestry, with an ancestry value to *icterina* of 0.744 and to *polyglotta* of 0.256. The difference in genotypes between both siblings could be explained either by a hybrid father or by two different fathers involved in the conception. As a support to this latter explanation, it has been shown that females in heterospecific pairs may engage in extra-pair copulations with conspecific males to reduce the costs of hybridization (Saetre & Saether 2010, Veen et al. 2001).

### Simulations

Based on the simulations, the applied set of microsatellites was suitable to detect hybrids between *Hippolais polyglotta* and *H. icterina*. The identified hybrids in this study were thus the first ones based on genetic evidence derived from microsatellites (*sensu* Engler et al. submitted). According to the simulations, individual DT12 was a F1 or later hybrid, whereas the other two individuals were likely backcrosses between a hybrid and a pure individual (*icterina* in the case of NL04 and *polyglotta* in DT06, Fig. 1a).

**Figure 1:**
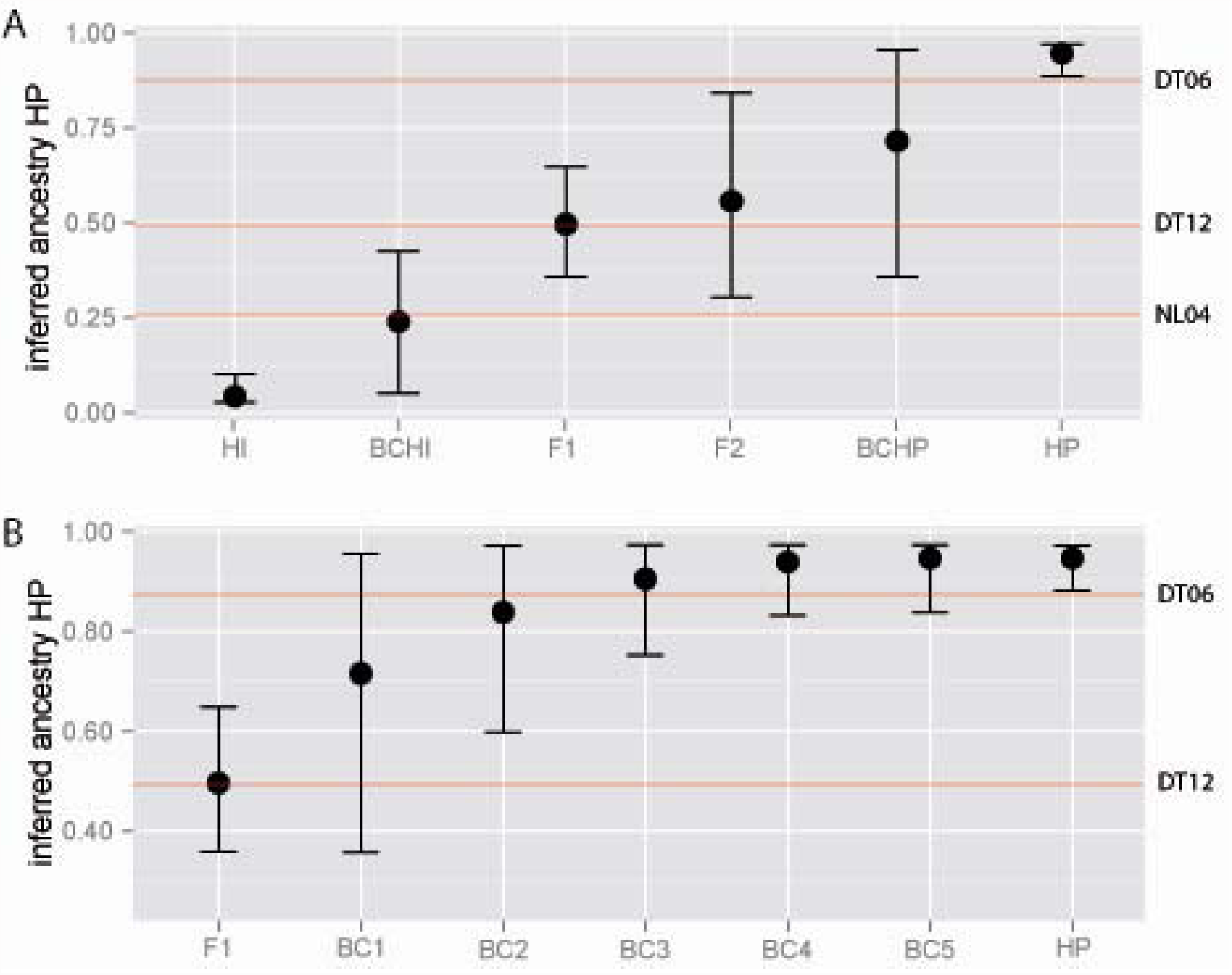
Ancestry inferred for hybrid simulated using allopatric genotypes of *Hippolais polyglotta* (HP) and *H. icterina* (HI) based on STRUCTURE. Shown are averages and respective 95% confidence limits. A) Direct hybrids (F1 & F2 generations) in comparison with backcrosses with either parental species (BCHP, BCHI) as well as with the parental species. B) Reduction of the genotypic signal of hybridization (F1) after five generations of backcrosses (BC1 – BC5) with the expansive species (HP). Red lines represent the inferred ancestry to *H. polyglotta* for the three hybrids detected (DT06, DT12, NL04).

The simulations further highlighted, that under conditions of permanent and unidirectional backcrossing (here from the receding *H. icterina* to the expansive *H. polyglotta*), drift quickly deletes hybrid alleles, supposedly neutral in SSR, (Fig 1b; Anderson & Thompson 2002). Thus, despite the use of a reasonably large set of microsatellites, the detection of introgression could be underestimated even if the true rate of hybridization (i.e. derived from pure parental individuals) is very accurate. In turn, focusing on AFLP markers seems to overestimate hybridization rate as discriminating between F1 hybrids and backcrosses is difficult in this dominant marker system. However, AFLP markers are able to track alleles longer due to a higher power, hence increasing the likelihood of detecting introgression. This is for two reasons: 1) generally much more loci are used in AFLP studies than in respective studies using SSR markers and 2) based on this, allele frequencies of alien alleles are higher given the larger number of loci to select from. Based on these stochastic differences between marker systems, erroneous estimation of hybridization and introgression rates could lead to wrong implications regarding the role and causes of hybridization in natural systems depending on the marker system used. Considering information provided by both marker systems, we can assume that even if the frequency of F1 hybrids is very low in these two *Hippolais* warblers, such individuals are viable and produce offspring (i.e. they generate backcrosses thus confirming the results from Secondi et al. 2006). However, after a few generations these backcrosses will be impossible to detect using SSR markers (Engler et al. unpublished).

This discrepancy in the results between the marker systems raises another issue about the role of adaptive responses during and after hybridization. In addition to the afore mentioned differences in power between SSR and AFLP markers, there are also differences regarding the neutrality of the marker system (ref). Fertile hybrids open the possibility to adaptive introgression, i.e. the transfer of beneficial alien genes (Arnold 2006, Kraus et al. 2012) into the gene-pool of the parental species. In moving contact zones, introgression can be highly biased towards the expanding species (Dasmahapatra et al. 2002, Secondi et al. 2006), as the density of the receding species is rapidly decreasing and homospecific partners of that species are in consequence less likely to find (Randler 2002). Therefore, unidirectional gene transfer could be expected in species pairs with moving contact zones and could be easier to detect in AFLP than in SSR marker systems.

In conclusion, hybridization quantified using neutral markers alone, such as microsatellites can give insights into the frequency of ongoing hybridization (i.e. by identifying F1 hybrids). Backcross-identification, however, strongly depends on the number of loci used and the frequency of species-specific alleles that these loci contain, with advantages of AFLP markers due to a higher power in detection of alien alleles. Single nucleotide polymorphisms (SNP) are a promising alternative in this regard as their high number lead to a high power in hybrid identification and thus provide a more accurate estimate of hybridization rate (references). They could also be used to search genes under selection. Therefore, attention has to be paid about the potential limits of classical marker systems to investigate the underlying processes involved in hybridization.

## Acknowledgements

We thank Boena van Noorden for sending us his Limburg samples and Jane Jönsson for aliquoting and sending JS’s samples from the storage in Lund to Trier. We are also grateful to Deborah A. Dawson and Joachim Kosuch who helped a lot with the selection of microsatellites as well as Petra Willems for genotyping the samples. This work is part of a larger study on the range expansion in H. polyglotta and would not have been possible without the outstanding contributions of Hilger Lemke. This project was kindly funded by the German Ornithological Society and by the Swiss Society for the Study and Conservation of Birds. JOE received additional funding from the Ministerium für Umwelt, Forsten und Verbraucherschutz Rheinland Pfalz and the German Federal Environmental Foundation. The authors declare no conflict of interests.

